# DeepCAGE: Incorporating Transcription Factors in Genome-wide Prediction of Chromatin Accessibility

**DOI:** 10.1101/610642

**Authors:** Qiao Liu, Kui Hua, Xuegong Zhang, Wing Hung Wong, Rui Jiang

## Abstract

Although computational approaches have been complementing high-throughput biological experiments for the identification of functional regions in the human genome, it remains a great challenge to systematically decipher interactions between transcription factors and regulatory elements to achieve interpretable annotations of chromatin accessibility across diverse cellular contexts. Towards this problem, we propose DeepCAGE, a deep learning framework that integrates sequence information and binding status of transcription factors, for the accurate prediction of chromatin accessible regions at a genome-wide scale in a variety of cell types. DeepCAGE takes advantage of a densely connected deep convolutional neural network architecture to automatically learn sequence signatures of known chromatin accessible regions, and then incorporates such features with expression levels and binding activities of human core transcription factors to predict novel chromatin accessible regions. In a series of systematic comparisons with existing methods, DeepCAGE exhibits superior performance in not only the classification but also the regression of chromatin accessibility signals. In detailed analysis of transcription factor activities, DeepCAGE successfully extracts novel binding motifs and measures the contribution of a transcription factor to the regulation with respect to a specific locus in a certain cell type. When applied to whole-genome sequencing data analysis, our method successfully prioritizes putative deleterious variants underlying a human complex trait, and thus provides insights into the understanding of disease-associated genetic variants. DeepCAGE can be downloaded from https://github.com/kimmo1019/DeepCAGE.

## Introduction

One of the fundamental questions in functional genomics is how activities of genes are spatially and temporally controlled through interactive effects of transcription factors (TFs) and regulatory elements (REs) such as promoters, enhancers and silencers. These regulatory elements, as short regions of non-coding DNA sequence, are known to typically reside in chromatin accessible regions and be bound by a set of TFs to carry out regulatory functionality in a manner specific to cellular contexts [1]. Therefore, the exploration of a landscape of chromatin accessible regions across major cell types will greatly facilitate the deciphering of gene regulatory mechanisms and further provide insights into cell differentiation, tissue homeostasis, and disease development [2].

Recent advances in deep sequencing techniques have enabled genome-wide assays of chromatin accessibility. For example, DNase-seq utilizes the DNase I enzyme to digest DNA sequences and identify DNase I-hypersensitive regions that are largely chromatin accessible [3]. ATAC-seq uses the Tn5 transposase to integrate primer DNA sequences into cleaved fragments that mainly come from chromatin accessible regions [4]. With the accomplishment of the ENCODE [5] and Roadmap [6] projects, these techniques have been successfully applied to the establishment of the chromatin accessibility landscape for dozens of cell lines across several species. The accumulation of these data provides an unprecedented opportunity for deepening our understanding of both gene regulation and occurrence of diseases [7,8,9].

However, due to limitations such as experimental cost, it is still impractical to further extend the landscape to cover all possible cell types, with the consideration of the huge variability in biological cellular contexts such as cell differentiation, environmental stimuli, and other factors. Towards this concern, computational approaches have been proposed to predict chromatin states by using such information as DNA sequence, gene expression, and other type of data [10–19]. For example, Kelley et al. proposed a deep convolutional neural network model called Basset to predict chromatin accessible regions purely relying on one-hot encoded DNA sequences [12]. Liu et al. developed a hybrid deep learning model for integrating multiple forms of sequence representations to achieve high prediction performance [14]. Quang et al. used a hybrid convolutional and recurrent neural network for predicting chromatin signals [18]. However, a model purely relying on sequence data can hardly be generalized to make predictions across different cell types as sequence itself is not cell-type-specific. To overcome this limitation, Zhou et al. proposed a regression model called BIRD that utilized only gene expression data to predict chromatin accessible regions [13]. Nevertheless, with the complete removal of sequence data, the scope of application of this method is limited, because the availability of gene expression is not as wide as sequence data. With the above understanding, Nair et al. proposed a deep residual neural network [20] model called ChromDragoNN to combine both sequence and expression data towards the prediction of chromatin accessibility [21]. However, sequence signatures and expression features are combined by simple concatenation in this method. This formulation, though simple in computation, lacks interpretability and is not consistent with existing biological knowledge.

With the above understanding, we propose a method called DeepCAGE, that is, a <u>**Deep**</u> densely connected convolutional network for predicting <u>**C**</u>hromatin <u>**A**</u>ccessibility by incorporating <u>**G**</u>ene <u>**E**</u>xpression and binding status of transcription factors. Unlike BIRD and ChromDragoNN that take full expression data as predictors, our method carefully considers the binding status of chromatin-binding factors (e.g., transcription factors), based on the biological understanding that chromatin accessibility is largely determined by chromatin-binding factors that have access to DNA [2]. In a series of systematic evaluation, DeepCAGE achieves state-of-the-art performance in not only the classification of chromatin accessible status, but also the regression of DNase-seq signals. To make DeepCAGE more understandable, we propose a strategy for visualizing the weights in the first convolutional layer. Interestingly, many known motifs were successfully recovered by DeepCAGE. In the downstream application to whole-genome sequencing (WGS) data analysis, DeepCAGE effectively prioritizes deleterious variants for the prediction and interpretation of complex phenotypes.

## Method

### Overview of DeepCAGE

DeepCAGE was designed based on the premise that binding status and gene expression of transcription factors could complement sequence data towards the precise prediction of chromatin accessibility. With this understanding, we designed DeepCAGE as a hybrid neural network that consisted of a convolutional module for sequence data and a feedforward module for chromatin accessibility prediction (**Figure 1**). Briefly, we applied the one-hot encoding to the input sequence data, fed the encoded data to a densely connected convolutional neural network (DenseNet), and took the output as the sequence feature. For binding status, we scanned the input sequence for potential binding sites for a set of 402 human transcription factors by using non-redundant motifs in the HOCOMOCO database [22] with the tool Homer [23]. We then selected the maximum score of reported binding sites for each transcription factor to obtain a vector of 402 dimensions as the motif feature. For gene expression, we focused on log-transformed TPM values of the 402 transcription factors and obtained a vector of 402 dimensions after quantile normalization as the expression feature. With these data, we combined the two vectors of motif and expression features by taking the element-wise product, and we concatenated the result to the sequence feature to obtain the hybrid feature, which went through a feedforward neural network with a fully connected hidden layer and an output layer for either classification or regression. We presented detailed hyperparameters of the hybrid network in (Table S1).

**Figure 1:**
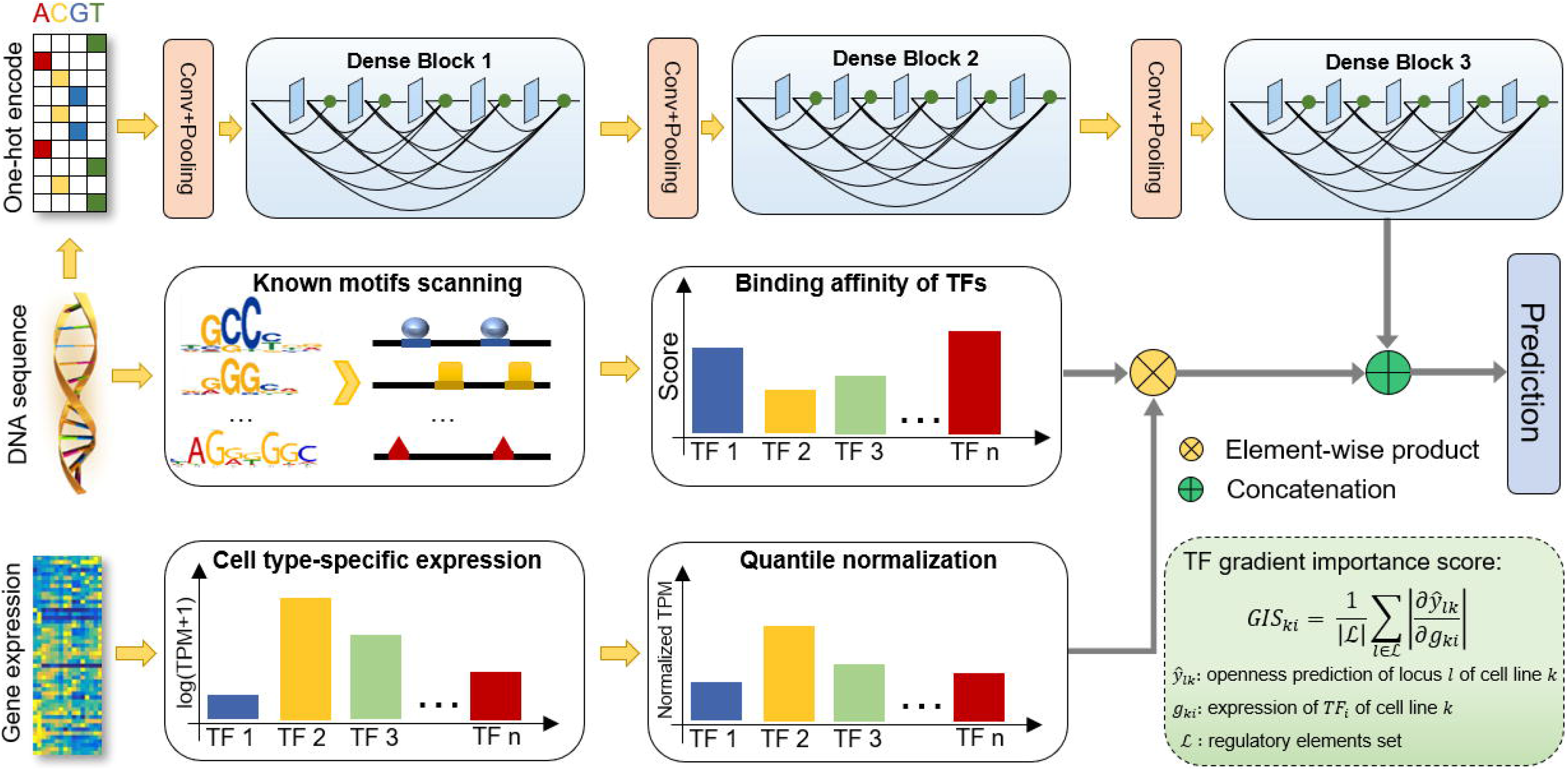
Overview of the DeepCAGE model. The sequence of the input DNA region is converted to a one-hot matrix and goes through a densely connected deep convolutional neural network (DenseNet) to extract sequence features. Normalized expression levels of the 402 human transcription factors and the corresponding motif binding scores are combined by using an element-wise product and then concatenated with sequence features. The combined features are finally fed to a feedforward neural network for chromatin accessibility prediction.

### Densely connected convolutional network

DeepCAGE extracts sequence features by using an architecture called the densely connected convolutional network (DenseNet), which has the advantage of alleviating the vanishing-gradient problem and strengthening the feature propagation [24]. As shown in Figure 1, there are three dense blocks in our model. Each block includes five convolutional layers, and each layer connects to every other layer in a feedforward fashion. A convolutional layer consists of two consecutive small kernels of size 1 × 1 and 3 × 1, where the former aims at reducing the concatenated channels to a fixed number, and the later acts as the traditional convolution. A transition module is presented before a dense block for feature extracting and dimensionality reduction. An input sequence is first extended to a fixed length of 1000 base pairs (bps) centered at the midpoint of the sequence and then converted to a 1000 × 4 binary matrix by using the one-hot encoding. The matrix is then fed to the first transition module that contains a convolutional layer and a max-pooling layer. The convolutional layer has 160 kernels of size 4 × 15 for extracting low-level features and detecting DNA binding motifs, while the max-pooling layer is present for finding the most significant activation signal in a given sliding window of each kernel. Similar settings are used for the other two transition modules for extracting high-level features and dimensionality reduction. Rectified linear units (ReLU) are used after each convolution operation for keeping positive activations and setting negative activation values to zeros. Batch normalization [25] and dropout [26] strategies are used after each ReLU function for reducing internal covariate shift and avoiding overfitting, respectively. For DeepCAGE regression model, there are two major differences from the classification model. First, the output layer directly uses a linear transformation instead of a sigmoid function. Second, the mean square error (MSE) instead of the cross-entropy is used as the loss function.

### Data processing

DNase-seq bam files and narrow peaks across 55 human cell types were downloaded from the ENCODE project [5] (Table S2-S3). The human hg19 reference genome was divided into non-overlapping regions (locus) of 200 bps. Considering that a cell type may have multiple DNase-seq replicates, a locus is regarded as chromatin accessible if it overlaps with narrow peak regions of at least half of the replicates and inaccessible otherwise (Figure S1). For the classification design, a binary label *y_lk_* is assigned to locus *l*, representing whether it is accessible in cell type *k*. For the regression design, bam files of multiple replicates for a cell type are pooled, and the raw read count, *n_lk_*, is obtained for locus *l* in cell type *k*. To eliminate the effect of sequencing depths, the normalized read count, 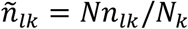, is calculated, where *N_k_* denotes the total number of pooled reads for cell type *k* and *N* = min{*N_k_*} the minimal number of pooled reads across all cell types. The normalized read counts are further log transformed after adding a pseudocount of one. The transformed data represent the level of chromatin accessibility and are then used as the response variable in the regression model.

RNA-seq data across the same 55 human cell types were also downloaded from the ENCODE project (Table S4). Gene expression data (TPM) of the 402 core human transcription factors were extracted. After further log transformation and quantile normalization based on TPM values, the normalized expression within each cell type was averaged across multiple replicates and the mean expression profile of each cell type was finally used.

Whole-genome sequencing (WGS) data and RNA-seq profiles of GTEx muscle tissues were downloaded from dbGaP (accession number phs000424.v7.p2). Matching these two types of data, a total of 491 donors were selected for downstream analysis (Table S5). For each of these donors, RNA-seq data was processed in the same way as ENCODE data, and WGS data was filtered by excluding all indels and rare variants whose minor allele frequencies were less than or equal to 5 across all donors.

### Model evaluation

Cell-type-level five-fold cross-validation experiments are designed for evaluating our method. In each fold, the 55 cell types are partitioned into a training set with 44 cell types and a test set with the remaining 11 cell types (Table S6-S7). Putative known accessible loci are identified as genomic regions (loci) that are chromatin accessible in at least two cell types in the training set. Putative novel accessible loci are identified as genomic regions that are accessible in at least two test cell types and are not present in the training data.

Cell-type-wise and locus-wise metrics are defined to evaluate our method from different perspectives (Figure S2). Cell-type-wise metrics are calculated within a test cell type across genomic regions to provide high level assessment of a method. Locus-wise metrics are calculated based on a genomic region across cell types to give detailed analysis of the performance. These metrics provide a comprehensive and systematic evaluation of our method in both the classification and the regression design.

Let ***Y***_*L*×*K*_ and 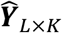 be the true label matrix and predicted matrix, where *L* denotes the number of putative loci and *K* that of cell types. In the classification design, *y_lk_* and 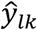 denote the true binary label and predicted probability of chromatin accessible status for locus *l* in cell type *k*, respectively. In this situation, the cell-type-wise auPR (area under the precision-recall curve) for cell type *k* is calculated based on ***y***_∗*k*_ = (*y*_1*k*_, *y*_2*k*_,…, *y*_*Lk*_) and 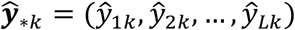 as follows. Given a threshold *t* for a cell type *k*, the precision is defined as the number of correct predictions 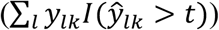 over the number of all predictions 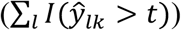, and the recall is defined as the number of correct predictions over the number of true accessible loci (∑_*l*_ *y_lk_*), where *I*(*x*) is an indicator function that is equal to 1 if *x* is true and 0 otherwise. Varying the threshold from 0 to 1 and calculating the precision and recall at each threshold value, the precision-recall curve can be drawn, and the area under this curve can be obtained. The locus-wise auPR for locus *l* is calculated based on ***y***_*l*∗_ = (*y*_*l*1_, *y*_*l*2_, *y*_*lk*_) and 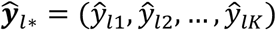 in a similar way.

In the regression design, *y_lk_* and 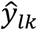 denote the true and predicted DNase-seq signal for locus *l* in cell type *k*, respectively. In this situation, the cell-type-wise PCC for cell type *k* is calculated as the Pearson correlation coefficient of ***y***_∗*k*_ and 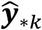, and the locus-wise PCC is calculated based on ***y***_*l*∗_ and 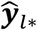 in a similar way. The prediction squared error (PSR), which considers both cell-type-wise prediction and locus-wise prediction, is calculated as 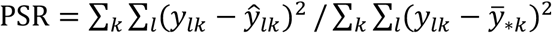, where 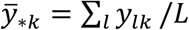 is the mean of ***y***_∗*k*_.

Two statistics, cell range and cell variability, are introduced to describe the activity of a locus based on the true DNase-seq signal across test cell types. The cell range of locus *l* is calculated by max(***y***_*l*∗_) − min (***y***_*l*∗_), and the cell variability of locus *l* is defined by the standard deviation of ***y***_*l*∗_.

### Baseline methods

Basset [12], DeepSEA [10], and DanQ [18] are three representative neural network models that take as input only DNA sequences. BIRD [13] is a regression model that takes as input only gene expression data. ChromDragoNN [21] is a neural network-based model that takes as input both DNA sequence and gene expression. Our method and ChromDragoNN have the following major differences. First, design principles of these two methods are notably different. ChromDragoNN predicts chromatin accessibility through direct concatenating of DNA sequence and expression data of all genes. DeepCAGE explains chromatin accessibility with DNA sequence and binding status of transcription factors. Therefore, DeepCAGE tries to interpret chromatin accessibility in a more natural way, since chromatin accessibility is believed to be largely determined by the occupancy and topological organization of nucleosomes as well as chromatin-binding factors [2]. Second, network architectures of these two methods are different. ChromDragoNN uses a ResNet to extract sequence features, while DeepCAGE uses a DenseNet that is a relatively new architecture and has also been experimentally validated to outperform ResNet in many tasks [20]. Third, inputs of these two methods are also different. ChromDragoNN requires DNA sequence and expression data of all genes, while DeepCAGE takes as input DNA sequence and expression data of 402 human core transcription factors. Motif binding profiles of these transcription factors can be annotated with existing motif database, which can be precomputed without additional experimental cost.

### Gradient importance score

DeepCAGE takes advantage of the gradient importance score (GIS) to prioritize transcription factors given a pair of cell type and a genomic locus. Briefly, a locus is extended to a 200kb genomic region centered at the midpoint of the locus. Then, the average absolute gradient of predicted accessibility within the extended region with respect to the expression of a transcription factor is calculated as

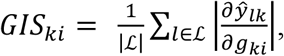

where, 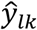 denotes the predicted accessibility of locus *l* in cell type *k*, *g_ki_* the expression of transcription factor *i* in cell type *k*, and 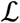 the set of putative regulatory elements that contains all accessible loci within the extended region. The gradient importance score gives an intuition of which transcription factors play an important role in a specific cell type.

### Motif analysis

The weights of the kernels from the first convolutional layer are converted into position weight matrices (PWMs) by counting subsequence occurrences in a set of input sequences that activate a kernel at a threshold value. All subsequences with activation value that greater than the threshold of a kernel are pooled together and aligned. The PWMs are then composed of the frequencies of the 4 nucleotides (A, C, G, and T) at each position. A subsequence at position *i* is regarded as activated if

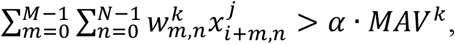

where *M* × *N* denotes the size of the kernels (4 × 15 in the first convolutional layer). *MAV^k^* denotes the maximal activation value of kernel *k* and is represented as

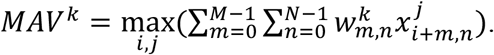

*α* is the control coefficient with the default value of 0.7 in all experiments.

Motifs are identified using the tool TomTom (v4.12.0) [27] with the E-value threshold 0.05 and are compared to known motifs in the JASPAR database (version 2018) [28]. Besides, the information content of recovered motifs is calculated based on the information entropy, as IC = ∑_*i,j*_(*p*_*i,j*_ log_2_ *p*_*i,j*_ − *b*_*i*_ log_2_ *b*_*i*_), where *p*_*i,j*_ is the element in PWM matrix, *i* and *j* the nucleotide type and position, respectively, and *b*_*i*_ (default value=0.25) the background frequency of nucleotide *i*.

### Phenotype prediction

A linear regression model with *l*_1_ penalty is adopted to predict the heights of GTEx donors using the deleterious scores of variants, as

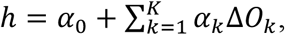

where *h* is the height of a GTEx donor, and Δ*O*_*k*_ denotes the deleterious score of variant *k* calculated using DeepCAGE. The coefficient of the *l*_1_ penalty is set to 0.5. A ten-fold cross validation experiment is used in validation, and the average coefficient of determinant (*R*^2^) is used for evaluating how much variance in the phenotype can be explained.

## Results

### DeepCAGE accurately predicts binary chromatin accessibility status

We first evaluated the performance of DeepCAGE in predicting whether an input DNA sequence is chromatin accessible or not. To achieve this objective, we downloaded paired DNase-seq and RNA-seq data across 55 cell types from the ENCODE project [5] and conducted a five-fold cross-validation experiment at the cell type level. In each fold of the validation, we partitioned the data into a training set of 44 cell types and a test set of the remaining 11 cell types. We then defined putative known accessible loci as genomic regions that are chromatin accessible in at least two cell types in the training data. For each cell type, we further identified a positive set as putative loci that were accessible in the cell type and a negative set as those inaccessible. After that, we trained our model on the training data and classified positive loci against negative ones for each test cell types. Finally, we calculated a criterion called the cell-type-wise auPR (see Methods) to evaluate the performance of a classification method.

We compared the performance of DeepCAGE with four existing methods, including Basset [12], DeepSEA [10], DanQ [18], and ChromDragoNN [21] in the above cross-validation experiment. Results (**Figure 2A**) show that DeepCAGE achieves the highest performance among all the methods with the mean cell-type-wise auPR of 0.418, compared to 0.166 of Basset, 0.195 of DeepSEA, 0.188 of DanQ, and 0.319 of ChromDragoNN. Particularly, DeepCAGE outperforms sequence-based methods by a large margin, suggesting that these methods may fail in capturing cell-type-specific information. Further analysis shows that the proportion of positive loci is in general small in a cell type and exhibits large variation (ranging from 2.6% to 29%), suggesting the ability of our method in dealing with unbalanced data.

**Figure 2:**
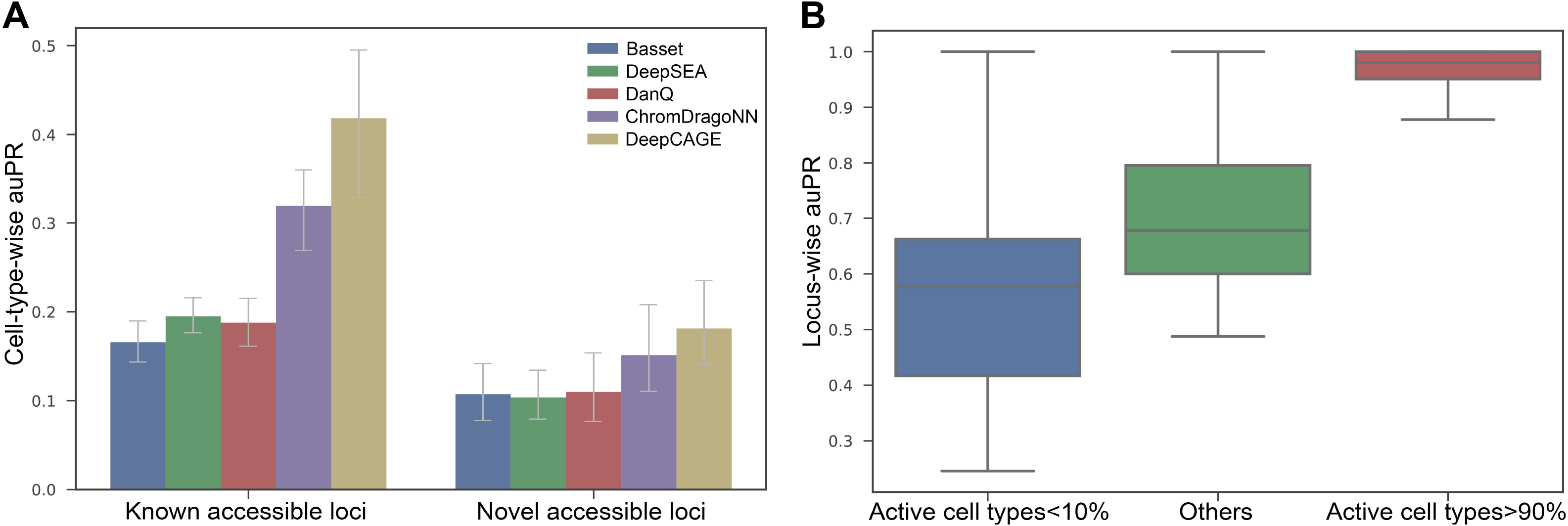
Performance of the DeepCAGE classification model. **A.**DeepCAGE achieves the highest cell-type-wise auPR for both known accessible loci and novel accessible loci compared to baseline methods (Basset, DeepSEA, DanQ and ChromDragoNN). **B.** The performance of DeepCAGE for loci with different activities across test cell types.

We then took one step further to access the ability of our method in predicting novel chromatin accessible loci. In each fold of the validation experiment, we identified putative novel accessible loci as genomic regions that were accessible in at least two test cell types and were not present in the training data, and we applied the trained model to predict whether these loci were accessible or not in a test cell type. Results, as shown in Figure 2A, also suggest the superiority of DeepCAGE with a mean cell-type-wise auPR of 0.181, compared to 0.107 of Basset, 0.104 of DeepSEA, 0.110 of DanQ, and 0.151 of ChromDragoNN.

We finally analyzed how the cell type specificity of accessible regions affects the prediction performance of our method. To achieve this objective, we divided the putative known accessible loci into three groups based on the proportion of cell types in which a locus is accessible. We then evaluated the cross-validation results using a criterion called the locus-wise auPR that evaluated the prediction performance of a method on an accessible locus across cell types (see Methods). Results show that for locus accessible in 10% cell types or less, DeepCAGE achieves a mean locus-wise auPR of 0.578, and this criterion increases when a locus is accessible in more cell types (Figure 2B). These results suggest that the cell type specificity is likely a factor that affects prediction performance of a method.

### DeepCAGE recovers continuous degree of chromatin accessibility

In the above classification experiments, we only consider the binary accessible status of a genomic region in a specific cell type. In the real situation, however, the accessibility of a genomic region given by a DNase-seq experiment is in continuous form. Considering this situation, we further proposed a DeepCAGE regression model to predict for a DNA region the degree of chromatin accessibility, which is defined as the normalized average count of raw reads that fall into the corresponding region.

With the same cross-validation settings as in the above section, we compared the performance of DeepCAGE to two baseline methods, BIRD [13] and ChromDragoNN [21], and we assessed regression results in term of two criteria, the cell-type-wise Pearson’s correlation coefficient (PCC) and prediction squared error (PSE) (see Methods). Results show that DeepCAGE achieves a mean cell-type-wise PCC of 0.785, compared to 0.637 for BIRD and 0.735 for ChromDragoNN (**Figure 3A**). Further analysis shows that in 18.2% of the test cell types, DeepCAGE achieves a cell-type-wise PCC of 0.85 or higher. In two cell types, DeepCAGE even achieves a cell-type-wise PCC of 0.9 or higher (see examples in Figure 3B). DeepCAGE also achieves the minimal PSE (0.420), outperforming the two baseline methods (0.770 for BIRD and 0.570 for ChromDragoNN) by a quite large margin (Figure 3C).

**Figure 3:**
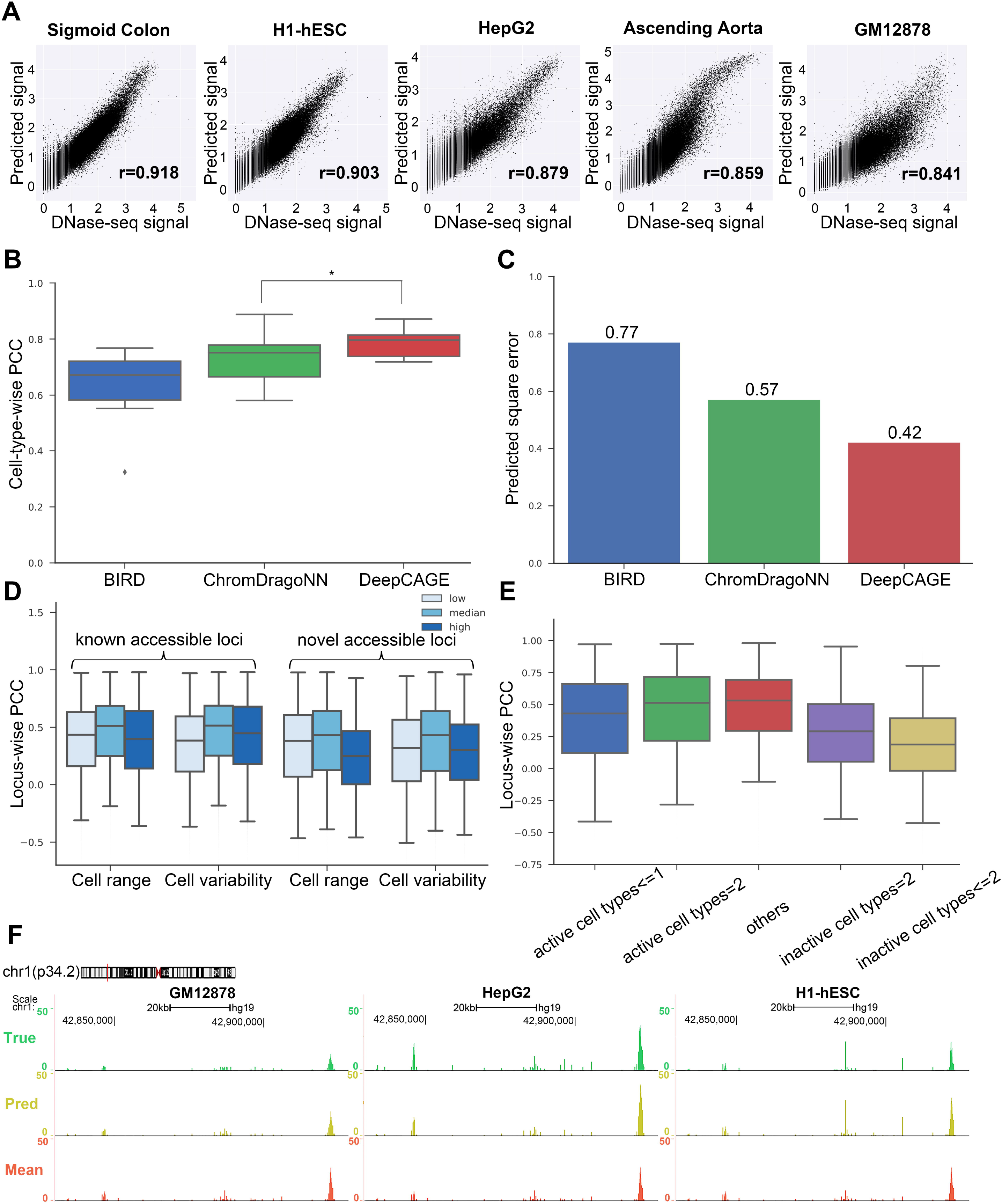
Performance of the DeepCAGE regression model. **A.** DeepCAGE predicts DNase-seq signals in five test cell types. **B.** Cell-type-wise PCC for three different methods across test cell types. * Two-sided paired-sample Wilcoxon signed-rank test *p*-value= 3.37 × 10^−5^. **C.** Prediction square error (PSR) for three different methods. **D.** Locus-wise PCC achieved by DeepCAGE with respect to two statistics with both known accessible loci and novel accessible loci. **E.** Locus-wise PCC achieved by DeepCAGE considering the number of accessible cell types under known accessible loci. **F.** An example of true (green) and predicted (yellow) DNase-seq signal of three test cell types under a same genomic region (chr1:42.83-42.93M). Mean signal (red) denotes the average DNase-seq signal across training cell types.

We then explored the performance of DeepCAGE for putative accessible loci with different cell type specificity by introducing two statistics, cell range and cell variability, to describe the activity dynamics of a genomic region based on the true DNase-seq signal cross cell types (see Methods). We divided known and novel accessible loci into three groups (low, median and high) according to the 1/3 and 2/3 quantiles of these statistics. Results show that DeepCAGE has high performance for accessible loci with medium cell range and cell variability (Figure 3D), consistent with the results in BIRD [13]. Briefly, DeepCAGE achieves a median locus-wise PCC (see Methods) of 0.512 for known accessible loci with medium cell range, compared to 0.435 and 0.399 for loci with low and high cell ranges, respectively. When using the statistic of cell variability, DeepCAGE achieves a median locus-wise PCC of 0.384, 0.514, and 0.448 for known accessible loci with low, median and high cell variabilities, respectively. The results are similar for novel accessible loci, except that values of the criteria are slightly low. We further divided known accessible loci into five groups based on the number of cell types in which a locus is accessible. Results (Figure 3E) show that the performance of DeepCAGE varies a lot for loci accessible in different numbers of cell types. Briefly, the performance is high for loci accessible in medium proportion of cell types and low for those accessible in only a small proportion of cell types.

Finally, we visualized in Figure 3F both the true (green) and predicted (yellow) DNase-seq signal of a sample genomic region across three test cell types (GM12878, HepG2, and H1-hESC) in the UCSC genome browser [29]. In contrast, we also provided as a reference the mean signal (red) that was calculated by taking the average DNase-seq signals across all training cell types. Obviously, DeepCAGE well distinguishes the difference of DNase-seq signals among the three test cell types while the mean signal fails.

### Model ablation analysis of DeepCAGE

We studied the contributions of gene expression and binding scores of transcription factors to the performance of our method. Taking the DeepCAGE regression model as an example, by discarding gene expression data, the median cell-type-wise PCC decreased by 13.1% (Figure S3, *p*-value=6.53×10^−11^ in a one-sided paired-sample Wilcoxon signed-rank test). By removing binding scores, the median cell-type-wise PCC decreased by 3.6% (Figure S3, *p*-value=3.78×10^−4^ in a one-sided paired-sample Wilcoxon signed-rank test). These results suggest that gene expression data could significantly help improve the performance of DeepCAGE in cross-cell-type prediction, while binding scores slightly increase the performance. One potential reason behind this observation is that a large proportion of DNA sequence motifs have already been learned in the convolution layers of the neural network, and thus the binding scores only provided complementary information regarding DNA sequence features.

Besides, to demonstrate the superiority of the network architecture used by DeepCAGE, we additionally conducted the following two experiments. First, we replaced the DenseNet with a ResNet which had the same number of layers as the number of dense blocks and the same hidden nodes in the convolutional layers. Results show that DenseNet leads to 6.4% increment in performance over ResNet in terms of the median cell-type-wise PCC (Figure S4, *p*-value=3.15×10^−6^ in a one-sided paired-sample Wilcoxon signed-rank test). Second, we explored the influence of two key hyperparameters (the number of residual blocks and the convolutional layers within a residual block) on the performance of ChromDragoNN. It is noted that a deeper model architecture does not help improve the performance significantly (Figure S5).

### Gradient importance score helps prioritize cell-type related transcription factors

We proposed a strategy for prioritizing cell-type related transcription factors according to the absolute gradient of the predicted accessibility with respect to the expression of a transcription factor. Taking the K562 cell line as an example, we calculated the average gradient importance score (GIS) of all transcription factors from all putative loci within up-streaming 100k bp to down-streaming 100k bp of a tumor suppressor gene *TP53*, which was shown to have a key role in myeloid blast transformation [30]. The average gradient importance score across cell types and transcription factors with respect to the transcription start site (TSS) of this gene was shown in **Figure 4A**. The 402 human core transcription factors were then prioritized by the average GIS score in K562 cell line (Figure 4B). Interestingly, many top-ranked transcription factors were related to functions in leukemia cell validated by literatures. For example, *EGR-*1 (rank^1st^) was involved in regulating PMA-induced megakaryocytic differentiation of K562 cell line [31]; the inhibition of *E2F7* (rank^3rd^) might lead to a reduction of miRNAs involved in leukemic cell lines [32]; the expression of *JunB* (rank^5th^) was inactivated by methylation in chronic myeloid leukemia [33]. The GO terms enriched by the top 5% prioritized transcription factor coding genes also included biological processes of leukocyte differentiation and hematopoietic development (Figure 4C). To sum up, the GIS score gives us an intuitive interpretation of which transcription factor may play an important role in predicting the chromatin accessibility given a specific cell type and a genomic region.

**Figure 4:**
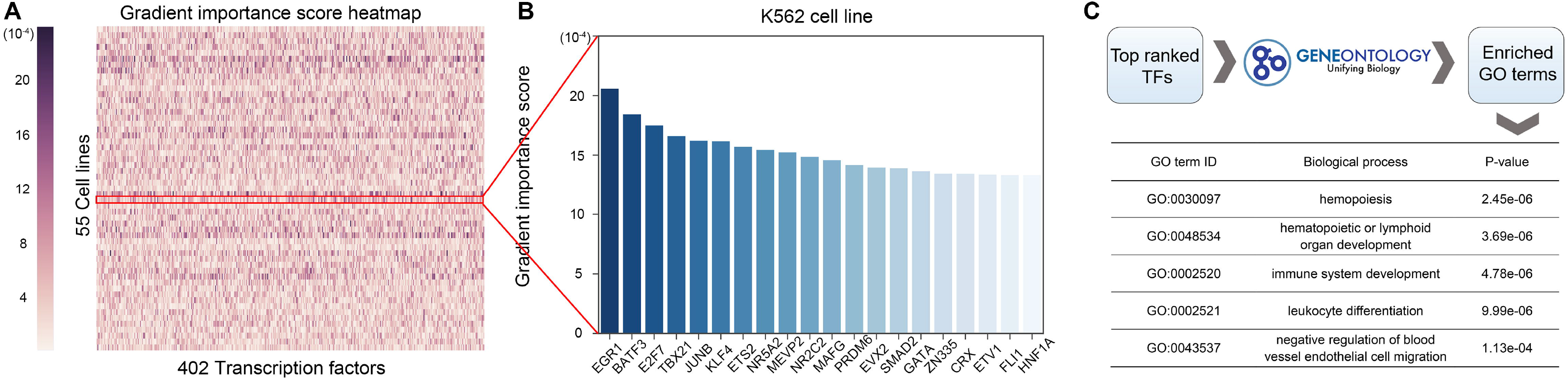
Gradient importance score helps identify important transcription factors. **A.** The GIS heatmap of the 402 human core transcription factors across 55 cell types. **B.** The gradient importance score of the 20 top-ranked transcription factors in the K562 cell type. **C.** Enriched GO terms by top-ranked transcription factors in the K562 cell type.

### DeepCAGE automatically learns binding motifs of transcription factors

In order to make DeepCAGE more understandable, we explored the features that were automatically learned by DeepCAGE by investigating the weights of the 160 kernels in the first convolutional layer. Briefly, we converted the weights into position weight matrices (PWMs) (see Methods) and then compared them with known motifs in the JASPAR database [28]. We found that 48 (30%) of the kernels could match known motifs at the E-value threshold of 0.05. Among the matched kernels, 25 (52%) had at least one matched core human transcription factor used in DeepCAGE model. We then calculated the information content (see Methods), set the weights of each kernel to zeros, and denoted the decrease in the cell-type-wise PCC as the influence score for each kernel. We showed several learned unmatched motifs that have a high influence score (**Figure 5A**) and illustrated a few examples of learned motifs that could match known motifs in JASPAR database (Figure 5B). These results demonstrate that DeepCAGE can not only help us find potential binding motifs, but also has the potential to guide the finding of novel motifs which are not discovered by experiments yet.

**Figure 5:**
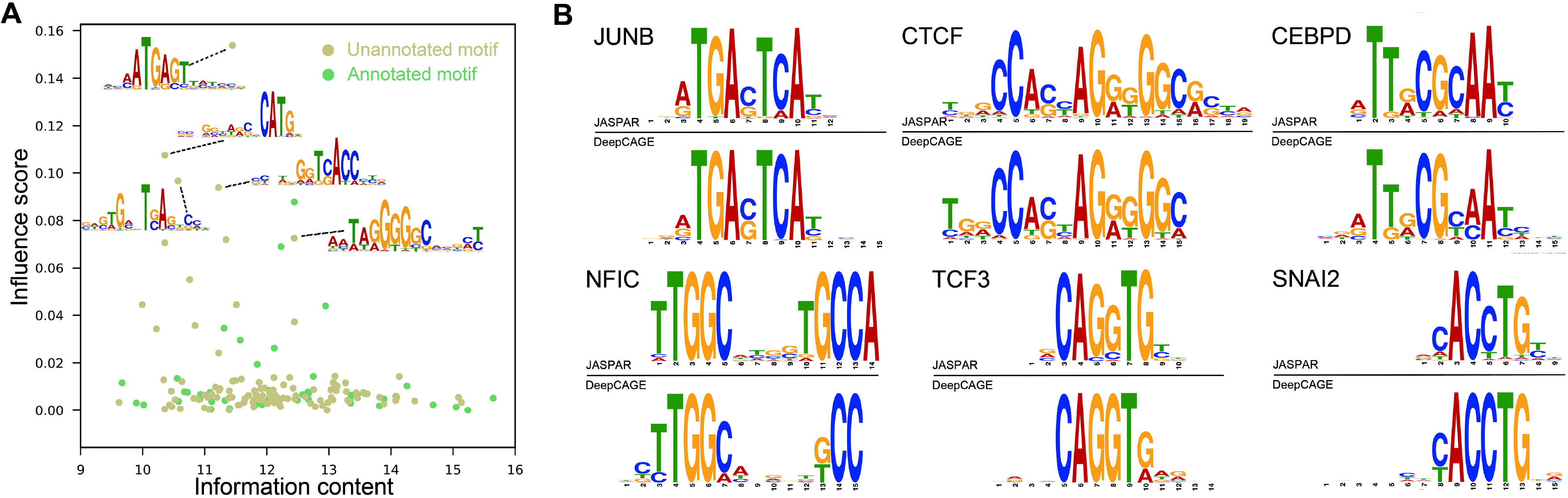
DeepCAGE recovers both known and novel motifs. **A.** Green dots and yellow dots represent known and novel motifs recovered by DeepCAGE, respectively. The x-axis describes the information content (see Methods) and the y-axis denotes the influence score. **B.** Matched motifs with an E-value threshold 0.05 in the format of sequence logos (above: known motif from the JASPAR database, below: motif learned by DeepCAGE).

### DeepCAGE prioritizes putative deleterious variants in WGS

We applied DeepCAGE to whole-genome sequencing (WGS) data analysis and demonstrated how our method could benefit the detection of individual-specific deleterious variants in regulatory elements that potentially influenced phenotype. The principle was to quantify the degree that a genetic variant affected the chromatin accessibility of a nearby genomic region and then prioritize variants accordingly. As shown in **Figure 6A**, for an individual, we fed the individual genome and the reference genome separately to the trained DeepCAGE regression model and calculated prediction scores for each of them. We then took the absolute log fold change of these two scores as a measure of the change in chromatin accessibility. For a variant, we defined its individual-level deleterious score by the change of chromatin accessibility of a 200bp genomic region around. Finally, we obtained the cohort-level deleterious score for a variant by applying the above procedure to all individuals in a cohort who contain the variant and then averaged individual-level deleterious scores for the variant.

**Figure 6:**
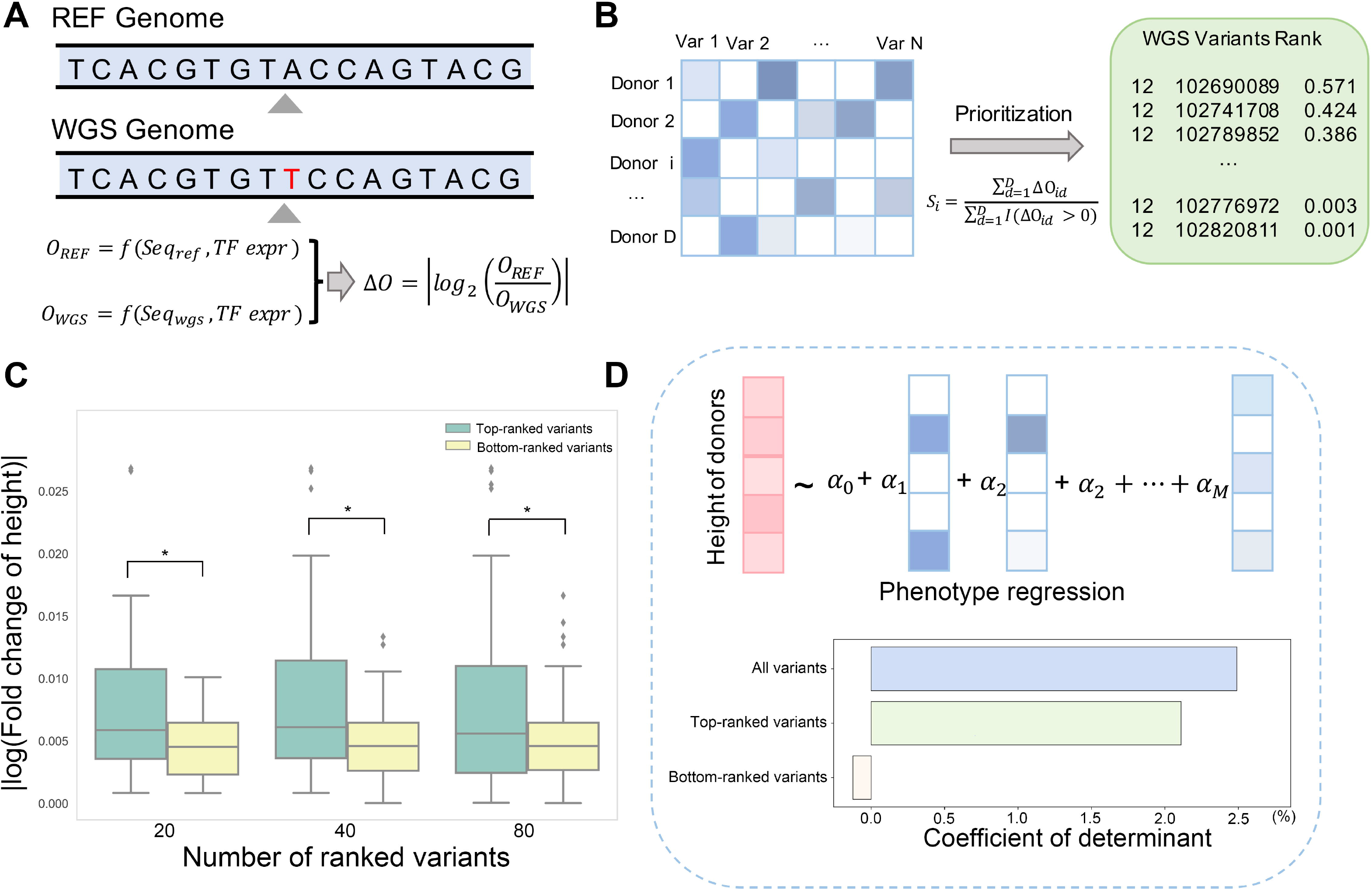
DeepCAGE helps prioritize and interpret WGS variants. **A.** The deleterious score is calculated by the absolute value of log change fold of predicted chromatin accessibility of the reference and WGS genomes. **B.** WGS variants within a risk region were ranked by averaging deleterious scores across donors containing the variant. **C.** The absolute log change fold of average height with respect to top-*K* and bottom-*K* ranked variants (*K*=20,40,80) around a height-associated gene. * indicates *p*-value<0.05. **D.** Predicting phenotype height with deleterious scores with all variants, top-ranked variants and bottom-ranked variants.

Note that we also took as input the expression profile of transcription factors in the muscle tissue and only consider WGS variant with the minor allele frequency larger than 5.

We downloaded WGS data of 491 donors with the height phenotype from the database of Genotypes and Phenotypes (dbGap) of GTEx project (Table S5). We collected 3290 risk single nucleic polymorphisms (SNPs) that were associated with height by a large-scale genome-wide association study [34]. For each risk SNP, we defined a risk region as a 200Kb genomic region centered at the SNP. We then ranked SNPs within a risk region according to their cohort-level deleterious scores obtained from donors (Figure 6B). As an illustration, we examined the risk region around a risk SNP (rs5742714) in the promoter region of *IGF1*, a well-known growth factor [35]. The top-ranked variants within this risk region showed an obviously higher absolute log fold change of average height than the bottom-ranked variants (Figure 6C).

We then quantitatively explored how much variance of the height phenotype can be explained by the deleterious scores of risk variants. To achieve this objective, we proposed a linear regression model with *l*_1_ penalty, which took deleterious scores of a set of variants as predictors and the height phenotype as the response variable (see methods). Results show that the 1,103,572 WGS variants within the 3290 risk regions together interpreted 2.49% of the height variance. Furthermore, the variants ranked among top 10% according to their deleterious scores in each risk region together can interpret 2.11% of the height variance. These results suggest that the small portion of variants prioritized by our method already contained most information that is helpful in the explanation of the phenotype. We also noticed that the bottom ranked 10% variants, on the contrary, failed to interpret the height phenotype (Figure 6D). To conclude, DeepCAGE is capable of giving a fine mapping of putative risk genetic variants and prioritizing WGS variants that might be associated with a specific phenotype.

## Discussion

In this paper, we have introduced a deep learning framework called DeepCAGE towards genome-wide prediction of chromatin accessibility. A hallmark of our method is the incorporation of sequence data and binding status of transcription factors into a unified deep neural network. With these two types of information complementing each other, our method overcomes limitations of existing approaches and demonstrates state-of-the-art performance in not only classification but also regression of chromatin accessibility signals. Our method provides insights into functional genomics in two aspects. First, the gradient importance score can give us an intuitional measurement of the contribution of a transcription factor to the regulation with respect to a specific locus in a certain cell type. Second, the visualization of convolutional kernels demonstrates that features automatically extracted by our method are not only consistent with existing knowledge, but also containing potentially novel binding motifs of transcription factors. Such interpretability of our model will benefit the dissection of regulatory landscape under a variety of cell conditions. Our method also provides the possibility of interpreting and prioritizing putative deleterious variants in genetic studies. Such ability in explaining complex traits can further be explored to promote the understanding of disease-associated genetic variants.

Certainly, our model can be further improved from the following aspects. First, currently we ignore the expression of genes that direct the synthesis of proteins other than transcription factors. However, it has been shown that proteins such as chromatin regulators, a class of enzymes with specialized function domains, can shape and maintain the epigenetic state in a cell context-dependent fashion [36], and thus can also provide information for inferring chromatin accessible state. How to incorporate information of these chromatin regulators into our model is one of the directions in our future work. Second, our model currently identifies chromatin accessible regions in a cell-type-specific manner, but cannot further distinguish the specific type of potential regulatory elements in these regions. With the accumulation of annotations regarding *cis*-regulatory elements such as enhancers [37] and silencers [38], as well as computational methods for predicting interactions between these elements [39–41], it is expected that our framework can further be extended to uncover the comprehensive relationship between different types of genomic regulatory elements and the genome-wide transcriptomic profile.

## Code availability

DeepCAGE is freely available at https://github.com/kimmo1019/DeepCAGE with step-by-step instructions.

## CRediT author statement

**Qiao Liu:** Conceptualization, Software, Formal analysis, Writing - Original Draft, Writing - Review & Editing, Visualization. **Kui Hua:** Writing - Review & Editing. **Xuegong Zhang:** Supervision. **Wing Hung Wong:** Conceptualization, Investigation, Supervision, Writing - Review & Editing, Funding acquisition. **Rui Jiang:** Conceptualization, Investigation, Supervision, Writing - Review & Editing, Funding acquisition. All authors read and approved the final manuscript.

## Acknowledgments

This work has been partly supported by the National Natural Science Foundation of China (Grant Nos. 61721003, 61873141, 61573207), the National Key R&D Program of China (Grant Nov. 2018YFC0910404), and the Tsinghua-Fuzhou Institute for Data Technology; This work was also supported by National Institutes of Health grants (Grant Nov. P50HG007735, R01HG010359). The Genotype-Tissue Expression project was supported by the Common Fund of the Office of the Director of the National Institutes of Health. We thank Mengmeng Wu, Zhana Duren and Fengling Chen for their helpful discussions.

## Supplementary material

**Figure S1 Identification of putative accessible loci**

Putative accessible loci were determined by DNase-seq peaks according to the following three steps. Step 1: The human reference genome was divided into non-overlapping regions of 200 bps. Step 2: Regions is kept if it is contained in half of replicated of a cell type. Step 3: Putative accessible loci were determined by collecting regions that appear in at least two cell types.

**Figure S2 Definition of cell type-wise and locus-wise criteria**

In the cell-type-wise evaluation, the area under the precision-recall curve (auPR) and Pearson’s correlation were calculated based on rows of both label matrix and predicted matrix. In the locus-wise evaluation, the Pearson’s correlation coefficient was calculated based on columns of both label matrix and predicted matrix.

**Figure S3 Model ablation analysis of DeepCAGE**

**A.** Four models were designed by considering different inputs. **B.** By removing both expression and motif scores of transcription factors (model 1), the median cell type-wise PCC decreases from 0.795 to 0.660. If only gene expression (model 2) or motif scores (model 3) are discarded, the median cell type-wise PCC decreases to 0.664 and 0.759, respectively.

**Figure S4 Ablation study for model architecture of DeepCAGE**

**A.** We implemented DeepCAGE model with two different architectures (DenseNet and ResNet). Note that we used a ResNet with three layers (equal to the number of dense blocks) and the number of hidden nodes and the convolutional kernel size is the same as convolutional layers in DenseNet. **B.** DeepCAGE with DenseNet architecture achieves a median cell-type-wise PCC of 0.795 while DeepCAGE with ResNet architecture achieves an average cell-type-wise PCC of 0.731.

**Figure S5 The cross-cell-type prediction performance of ChromDragoNN with different hyperparameter settings**

**A.** The cell-type-wise PCC of ChromDragoNN with different number of convolutional layers in each residual block. **B.** The cell-type-wise PCC of ChromDragoNN with different number of residual blocks.

**Table S1 Hyperparameters of the DeepCAGE model**

**Table S2 The information of Dnase-seq peaks across 55 cell types from the ENCODE project**

**Table S3 The information of Dnase-seq bam file across 55 cell types from the ENCODE project**

**Table S4 The information of RNA-seq data across 55 cell types from the ENCODE project**

**Table S5 The information of GTEx data collected from 491 donors**

**Table S6 Information of the 55 cell types used in this study**

**Table S7 Data partition in the five-fold cross-validation experiment**

## References

[1] Kellis M, Wold B, Snyder MP, Bernstein BE, Kundaje A, Marinov GK, et al. Defining functional DNA elements in the human genome. Proc Natl Acad Sci USA 2014;111:6131–8.

[2] Klemm SL, Shipony Z, Greenleaf WJ. Chromatin accessibility and the regulatory epigenome. Nat Rev Genet 2019;20:207–20.

[3] Crawford GE, Holt IE, Whittle J, Webb BD, Tai D, Davis S, et al. Genome-wide mapping of DNase hypersensitive sites using massively parallel signature sequencing (MPSS). Genome Res 2006;16:123–31.

[4] Buenrostro JD, Giresi PG, Zaba LC, Chang HY, Greenleaf WJ. Transposition of native chromatin for fast and sensitive epigenomic profiling of open chromatin, DNA-binding proteins and nucleosome position. Nat Methods 2013;10:1213.

[5] Consortium EP. An integrated encyclopedia of DNA elements in the human genome. Nature 2012;489:57–74.

[6] Roadmap Epigenomics C, Kundaje A, Meuleman W, Ernst J, Bilenky M, Yen A, et al. Integrative analysis of 111 reference human epigenomes. Nature 2015;518:317–30.

[7] Corces MR, Granja JM, Shams S, Louie BH, Seoane JA, Zhou W, et al. The chromatin accessibility landscape of primary human cancers. Science 2018;362.

[8] Trevino AE, Sinnott-Armstrong N, Andersen J, Yoon S-J, Huber N, Pritchard JK, et al. Chromatin accessibility dynamics in a model of human forebrain development. Science 2020;367.

[9] Song S, Cui H, Chen S, Liu Q, Jiang R. EpiFIT: functional interpretation of transcription factors based on combination of sequence and epigenetic information. Quant Biol 2019;7:233–43.

[10] Zhou J, Troyanskaya OG. Predicting effects of noncoding variants with deep learning-based sequence model. Nat Methods 2015;12:931–4.

[11] Liu Q, Gan M, Jiang R. A sequence-based method to predict the impact of regulatory variants using random forest. BMC Syst Biol 2017;11:7.

[12] Kelley DR, Snoek J, Rinn JL. Basset: learning the regulatory code of the accessible genome with deep convolutional neural networks. Genome Res 2016;26:990–9.

[13] Zhou W, Sherwood B, Ji Z, Xue Y, Du F, Bai J, et al. Genome-wide prediction of DNase I hypersensitivity using gene expression. Nat Commun 2017;8:1038.

[14] Liu Q, Xia F, Yin Q, Jiang R. Chromatin accessibility prediction via a hybrid deep convolutional neural network. Bioinformatics 2018;34:732–8.

[15] Min X, Zeng W, Chen N, Chen T, Jiang R. Chromatin accessibility prediction via convolutional long short-term memory networks with k-mer embedding. Bioinformatics 2017;33:i92–i101.

[16] Xu C, Liu Q, Zhou J, Xie M, Feng J, Jiang T. Quantifying functional impact of non-coding variants with multi-task Bayesian neural network. Bioinformatics 2020;36:1397–404.

[17] Yin Q, Wu M, Liu Q, Lv H, Jiang R. DeepHistone: a deep learning approach to predicting histone modifications. BMC genomics 2019;20:11–23.

[18] Quang D, Xie X. DanQ: a hybrid convolutional and recurrent deep neural network for quantifying the function of DNA sequences. Nucleic Acids Res 2016;44:e107–e.

[19] Ding K, Liu Q, Lee E, Zhou M, Lu A, Zhang S. Feature-Enhanced Graph Networks for Genetic Mutational Prediction Using Histopathological Images in Colon Cancer. International Conference on Medical Image Computing and Computer-Assisted Intervention 2020:294–304.

[20] He K, Zhang X, Ren S, Sun J. Deep residual learning for image recognition. Proceedings of the IEEE conference on computer vision and pattern recognition 2016:770–8.

[21] Nair S, Kim DS, Perricone J, Kundaje A. Integrating regulatory DNA sequence and gene expression to predict genome-wide chromatin accessibility across cellular contexts. Bioinformatics 2019;35:i108–i16.

[22] Kulakovskiy IV, Vorontsov IE, Yevshin IS, Soboleva AV, Kasianov AS, Ashoor H, et al. HOCOMOCO: expansion and enhancement of the collection of transcription factor binding sites models. Nucleic Acids Res 2015;44:D116–D25.

[23] Heinz S, Benner C, Spann N, Bertolino E, Lin YC, Laslo P, et al. Simple combinations of lineage-determining transcription factors prime cis-regulatory elements required for macrophage and B cell identities. Molecular cell 2010;38:576–89.

[24] Huang G, Liu Z, Weinberger KQ, van der Maaten L. Densely connected convolutional networks. Proceedings of the IEEE conference on computer vision and pattern recognition 2017;1:3.

[25] Ioffe S, Szegedy C (2015), ‘Batch Normalization: Accelerating Deep Network Training by Reducing Internal Covariate Shift’, in Francis B., David B. Eds., Proceedings of the 32nd International Conference on Machine Learning, PMLR, Proceedings of Machine Learning Research, pp. 448–56.

[26] Srivastava N, Hinton G, Krizhevsky A, Sutskever I, Salakhutdinov R. Dropout: A simple way to prevent neural networks from overfitting. The Journal of Machine Learning Research 2014;15:1929–58.

[27] Gupta S, Stamatoyannopoulos JA, Bailey TL, Noble WS. Quantifying similarity between motifs. Genome Biol 2007;8:R24.

[28] Khan A, Fornes O, Stigliani A, Gheorghe M, Castro-Mondragon JA, van der Lee R, et al. JASPAR 2018: update of the open-access database of transcription factor binding profiles and its web framework. Nucleic Acids Res 2017;46:D260–D6.

[29] Kent WJ, Sugnet CW, Furey TS, Roskin KM, Pringle TH, Zahler AM, et al. The human genome browser at UCSC. Genome Res 2002;12:996–1006.

[30] Law JC, Ritke MK, Yalowich JC, Leder GH, Ferrell RE. Mutational inactivation of the p53 gene in the human erythroid leukemic K562 cell line. Leukemia research 1993;17:1045–50.

[31] Cheng T, Wang Y, Dai W. Transcription factor egr-1 is involved in phorbol 12-myristate 13-acetate-induced megakaryocytic differentiation of K562 cells. J Biol Chem 1994;269:30848–53.

[32] Gabra MM, Salmena L. microRNAs and Acute Myeloid Leukemia chemoresistance: a mechanistic overview. Frontiers in oncology 2017;7.

[33] Yang M-Y, Liu T-C, Chang J-G, Lin P-M, Lin S-F. JunB gene expression is inactivated by methylation in chronic myeloid leukemia. Blood 2003;101:3205–11.

[34] Yengo L, Sidorenko J, Kemper KE, Zheng Z, Wood AR, Weedon MN, et al. Meta-analysis of genome-wide association studies for height and body mass index in∼ 700000 individuals of European ancestry. Hum Mol Genet 2018;27:3641–9.

[35] Becker NS, Verdu P, Georges M, Duquesnoy P, Froment A, Amselem S, et al. The role of GHR and IGF1 genes in the genetic determination of African pygmies’ short stature. Eur J Hum Genet 2013;21:653–8.

[36] Chen T, Dent SY. Chromatin modifiers and remodellers: regulators of cellular differentiation. Nat Rev Genet 2014;15:93.

[37] Khan A, Zhang X. dbSUPER: a database of super-enhancers in mouse and human genome. Nucleic Acids Res 2016;44:D164–D71.

[38] Zeng W, Chen S, Cui X, Chen X, Gao Z, Jiang R. SilencerDB: a comprehensive database of silencers. Nucleic Acids Res 2021;49:D221–D8.

[39] Li W, Wong WH, Jiang R. DeepTACT: predicting 3D chromatin contacts via bootstrapping deep learning. Nucleic Acids Res 2019;47:e60–e.

[40] Liu Q, Lv H, Jiang R. hicGAN infers super resolution Hi-C data with generative adversarial networks. Bioinformatics 2019;35:i99–i107.

[41] Singh S, Yang Y, Poczos B, Ma J. Predicting enhancer-promoter interaction from genomic sequence with deep neural networks. Quant Bio 2019;7:122–37.

